# Target-detecting neurons in the dragonfly ‘lock on’ to selectively attended targets

**DOI:** 10.1101/496562

**Authors:** B.H. Lancer, B.J.E. Evans, J.M. Fabian, D.C. O’Carroll, S.D. Wiederman

## Abstract

The visual world projects a complex and rapidly changing image on to the retina, presenting a computational challenge for any animal relying on vision for an accurate view of the world. One such challenge is parsing a visual scene for the most salient targets, such as the selection of prey amidst a swarm. The ability to selectivity prioritize processing of some stimuli over others is known as ‘*selective attention’*. Previously, we identified a dragonfly visual neuron called ‘Centrifugal Small Target Motion Detector 1’ (CSTMD1) that exhibits selective attention when presented with multiple, equally salient features. Here we conducted electrophysiological recordings from CSTMD1 neurons *in vivo*, whilst presenting visual stimuli on a monitor display. To identify the target selected in any given trial, we modulated the intensity of moving targets, each with a unique frequency (frequency-tagging). We find that the frequency information of the *selected* stimulus is preserved in the neuronal response, whilst the distracter is completely ignored. We show that the competitive system that underlies selection in this neuron can be biased by the presentation of a preceding target on the same trajectory, even when it is of lower contrast to the distracter. With an improved method of identifying and biasing target selection in CSTMD1, the dragonfly provides an effective animal model system to probe the mechanisms underlying neuronal selective attention.

**Significance Statement:** This is a novel application of frequency tagging at the intracellular level, demonstrating that frequency information of a flickering stimulus is preserved in the response of an individual neuron. Using this technique, we show that the selective attention mechanism in an individual dragonfly visual neuron is able to lock on to the selected stimuli, in the presence of distracters, even those of abrupt onset or higher contrast. Conversely, unidentified factors allow selection to occasionally switch mid-trial to the other target. We therefore show that this neuronal network underlying selective attention is more complex than the traditionally modelled winner-takes-all framework.

## Introduction

The visual world contains a wealth of information about the environment and surroundings, but even the most sophisticated visual systems lack the capacity to encode all the information contained in a scene over time. Instead, animals must parse a scene for behaviourally relevant information and discard the remaining clutter. One solution to this problem is *selective attention*, the ability to selectively respond to one stimulus amongst multiple alternatives. Selective attention is observed across species, from humans and other primates (Treue, 2001), to ‘simple’ insects, including the fruit fly (De Bivort & van Swinderen, 2016; Nityananda 2016). Selective attention is particularly important in predatory animals which hunt among swarms containing potentially hundreds of prey and conspecifics, such as the dragonfly (Edman & Haeger, 1974; Baird & May, 1997). Many predators hunting in these conditions are susceptible to the ‘confusion effect’, a reduced success rate due to difficulty tracking a single target amidst the swarm (Landeau & Terborgh, 1986, Jeschke & Tollrian, 2007). Some dragonflies, however, show particularly good performance hunting among swarms across all stages of life (Jeschke & Tollrian, 2007; Combes et al, 2012).

Successful prey capture relies on the ability to filter irrelevant information, such as background clutter and conspecifics, whilst selecting and tracking prey amongst equally valuable alternatives. Indeed, the confusion effect is diminished where predators are able to visually identify and track individual prey (Landeau & Terborgh, 1986). In order to achieve this, the underlying neuronal system must be able to ‘lock-on’ to and track an individual, noisy target, while simultaneously flexible enough to switch targets when this would increase the chance of success.

We have previously identified a visual neuron in the dragonfly optic lobe that exhibits a ‘winner-takes-all’ selective attention (Wiederman & O’Carroll, 2013), named ‘Centrifugal Small-Target Motion Detector’ (CSTMD1). CSTMD1 is tuned for the movement of small (1°- 3°) dark targets against a bright background (O’Carroll 1993; Geurten et al, 2007), matching the demands of an ethologically relevant target-detection system (Labhart & Nilsson, 1994; Olberg et al, 2005; Olberg et al., 2007). When presented with two such targets, CSTMD1 encodes the *absolute* strength of the selected target without interference from distracters (Wiedermen & O’Carroll, 2013). In contrast, typical findings in primates (eg. Recanzone, Wurtz & Schwarz, 1997; Treue & Maunsell, 1999), Owls (Asadollahi, Mysore & Knudsen, 2010) and other insects (Tang & Juusola, 2010; van Swinderen, 2012) show a response that is modulated by the presence of non-selected distracters. Encoding an absolute representation of a selected target (i.e. ignoring the distracter) has been observed in the auditory system of crickets (Pollack, 1998) and in primate neurons in MT (Harrison, Weiner & Ghose, 2013). The analogue of CSTMD1 processing in human psychophysics is ‘inattentional blindness’, whereby an object in the visual field is ignored while attention is focused elsewhere (Simons & Chabris, 1999).

Previously, we have shown that CSTMD1 exhibits properties important for a prey-tracking system. Firstly, the rare observation that selection could switch between targets mid-way through a trial (Wiederman & O’Carroll, 2013). This raised the intriguing possibility that an ongoing competitive mechanism drives target selection, even after an initial target has been selected, and that this mechanism can direct switches at opportune moments. Secondly, CSTMD1 exhibits ‘predictive gain modulation’ whereby a local faciltatory ‘spotlight’ of increased gain spreads forward along the predicted trajectory of a target (even accounting for occlusions), with inhibition elsewhere in the receptive field surround (Dunbier et al, 2012; Wiederman, Fabian et al., 2017). This facilitation may represent a mechanism for ‘locking-on’ to a selected target, for example, a chosen fruitfly in a swarm.

Here, we have developed a technique to frequency-tag targets by exploiting their contrast dependant response (O’Carroll & Wiederman, 2014), thus permitting us to determine which target has been selected at any moment. We show that CSTMD1 is both able to dynamically switch selected targets mid-trail and lock-on to selected targets, even in the presence of a higher contrast distracter. We therefore describe a neuronal system more complex than the traditionally modelled winner-takes-all framework. This provides important insight into how selective behaviours are implemented by underlying neuronal processing.

## Materials & Methods

### Experimental Preparation

We recorded from a total of 26 male, wild-caught *Hemicordulia tau* dragonflies. Dragonflies were stored at 7°c for up to 7 days before experimentation. Dragonflies were warmed and then immobilized to an articulating magnetic stand with a 50/50 wax-rosin mixture. The head was tilted forwards to allow access to the back of the head, and a small hole was dissected in the rear of the head capsule adjacent to the oesophagus to allow visual and physical access to the brain.

We pulled aluminosilicate electrodes (Harvard Apparatus) using a Sutter Instruments P-97 electrode puller, which were filled with a 2M KCl solution. Electrodes were then inserted into the lobula complex using a piezo-electric stepper with a typical resistance of 40-140 MΩ. Intracellular responses were digitised at 5 kHz for offline analysis with MATLAB.

### Visual Stimuli

We presented stimuli on high-definition LCD computer monitors (120 – 165 Hz) using a custom-built presentation and data acquisition suite based on MATLAB (RRID: SCR_001622) and PsychToolBox (RRID: SCR_002881. Available: http://psychtoolbox.org/). The animal was placed 20 cm away from the monitor and centred on the visual midline, thus minimizing off-axis artefacts. Stimuli consisted of a single or pair (∼20° separation) of 1.5° by 1.5° squares of modulated contrast ascending the receptive field at a speed of 40°/s.

We applied to our intracellular recordings a frequency-tagging paradigm inspired by human electroencephalography research (Norcia et al., 2015) and local field potential research in insects (van Swinderen, 2012). We presented two competing, flickering targets each with varying contrast at two different frequencies. As neuronal responses are themselves modulated by the contrast, the response frequency permits identification of the selected target. We presented non-harmonic frequency-pairs of either 8 Hz (*F*_*1*_) and 12 (*F*_*2*_) Hz, or 11 Hz (*F*_*1*_) & 15(*F*_*2*_) Hz. The two frequency pairs tested the robustness of the technique as well as ensuring that there was no artefacts induced from interactions with the display refresh rates. That is, the frequencies were not multiples of one other and were divisible by the monitor refresh rate thus ensuring the full range of intensities were presented within each period. We tested with both sinusoidal and square wave flicker. These results were pooled because there was no difference in their power to identify selection.

Frequency tagged targets flickered between a minimum Weber contrast of 0.06 and maximum of 1 (mean contrast of 0.51 and a white background of 337 Cd/m^2^). In single target trials, one target contrast varied at either F_1_, F_2_, or 0 Hz (Non-flickering control at maximum contrast) and was presented moving vertically up the display at one of two spatial locations, T_1_ or T_2_ (locations separated 20° horizontally within CSTMD1’s receptive field). In paired target trials, two flickering targets were presented at T_1_ and T_2_ locations. The choice whether the spatial location T_1_ or T_2_ was either F_1_ or F_2_ (e.g. 8 Hz or 12 Hz), was pseudo-randomized to control for any preferred frequency response.

### Experimental design and statistical analysis

For testing hypotheses about trial by trial selection processes, any given trial is an independent event and cannot be averaged as a technical replicate. However, to ensure robustness of the result we repeated experiments across a number of dragonflies. Here we use ‘n’ to denote the number of trials and additionally report across how many dragonflies. We visualise all trial data points and describe similarities or differences across animals.

We report exact P except when less than 0.001. All tests are nonparametric, two-tailed and corrected for multiple comparisons (Bonferroni-Holm correction). Box & Whisker plots indicate median, interquartile and minimum/maximum range. Unless otherwise stated outliers are indicated with crosses.

All data analysis was conducted in MATLAB 2017a (RRID: SCR_001622), including the Wavelet Toolbox. Complete Wavelet Transforms (CWT’s) used an analytic Morse wavelet with gamma = 3.

## Results

### Neuronal responses are Frequency Tagged

To test the validity of the frequency tagging technique, we presented a single flickering target moving vertically up the display within the dragonfly’s field of view (**Figure 1A**). The target drifted at 40°/s within the excitatory, contralateral region of CSTMD1’s receptive field (Wiederman and O’Carroll, 2013; Wiederman, Fabian et al, 2017). We use the term ‘Frequency tagging’ to refer to the modulation of Weber contrast: (Intensity_target_ - Intensity_background_) / I_background_, over time at a set frequency (in Hertz). Since CSTMD is responsive to dark targets (Wiederman, Shoemaker & O’Carroll, 2013), we flickered a black-to-grey target against a white background (**Figure 1B**). An example of an individual data trace in response to a 15 Hz target shows the spike activity during the stimulus presentation (**Figure 1C**, dark bar). To extract any frequency-tagged response modulation, we first determine spike locations and calculate the instantaneous spike rate (Inverse Inter-Spike Interval) over time (**Figure 1D**). We then apply one of two mathematical transforms to this data. The application of a Fast Fourier Transform (square root to provide amplitude) reveals a peak in the frequency domain at 15 Hz (**Figure 1E**), equivalent to the target contrast modulation (a response at 0 Hz is due to the non-zero mean over time). We repeated this process for a series of different frequencies (averaged across neurons) to determine the most appropriate for further experiments (**Figure 1F**). This data shows that from 7 to 19 Hz the frequency content of the stimuli is well preserved in the intracellular response of single neurons. However, we have previously shown that CSTMD1 can ‘switch’ selection mid-trial (Wiederman & O’Carroll, 2013). In this circumstance, power in an FFT would be distributed between the two target frequencies, corresponding to the total time each target was selected. Therefore, Fourier analysis cannot distinguish when: (1) trials where modulation was genuinely shared between T_1_ and T_2_ (indicative of a lack of competitive selection, such as neuronal summation) or (2) selection switched from T_1_ to T_2_ or T_2_ to T_1_ mid-way through the trial. To account for possible switches, we instead applied Continuous Wavelet Transforms (CWTs) which provide an approximate power across pseudo-frequencies *over time*. Averaging this wavelet analysis across time is similar to an FFT though reveals a broader peak in the frequency domain centred at 15 Hz (**Figure 1G**). The broader shape observed in the CWT is inherent to the wavelet analysis, and is the cost of providing information of how frequency components might vary over time. Although in the frequency domain CWT responses are blurred in comparison to their FFT counterparts, there are statistically significant differences for any two frequencies separated by at least 2 Hz (*P <.001*). Thus we were able to analyse all further data using CWTs to derive the benefit of examining the frequency response evolution over time of the individual trials.

**Figure 1:**
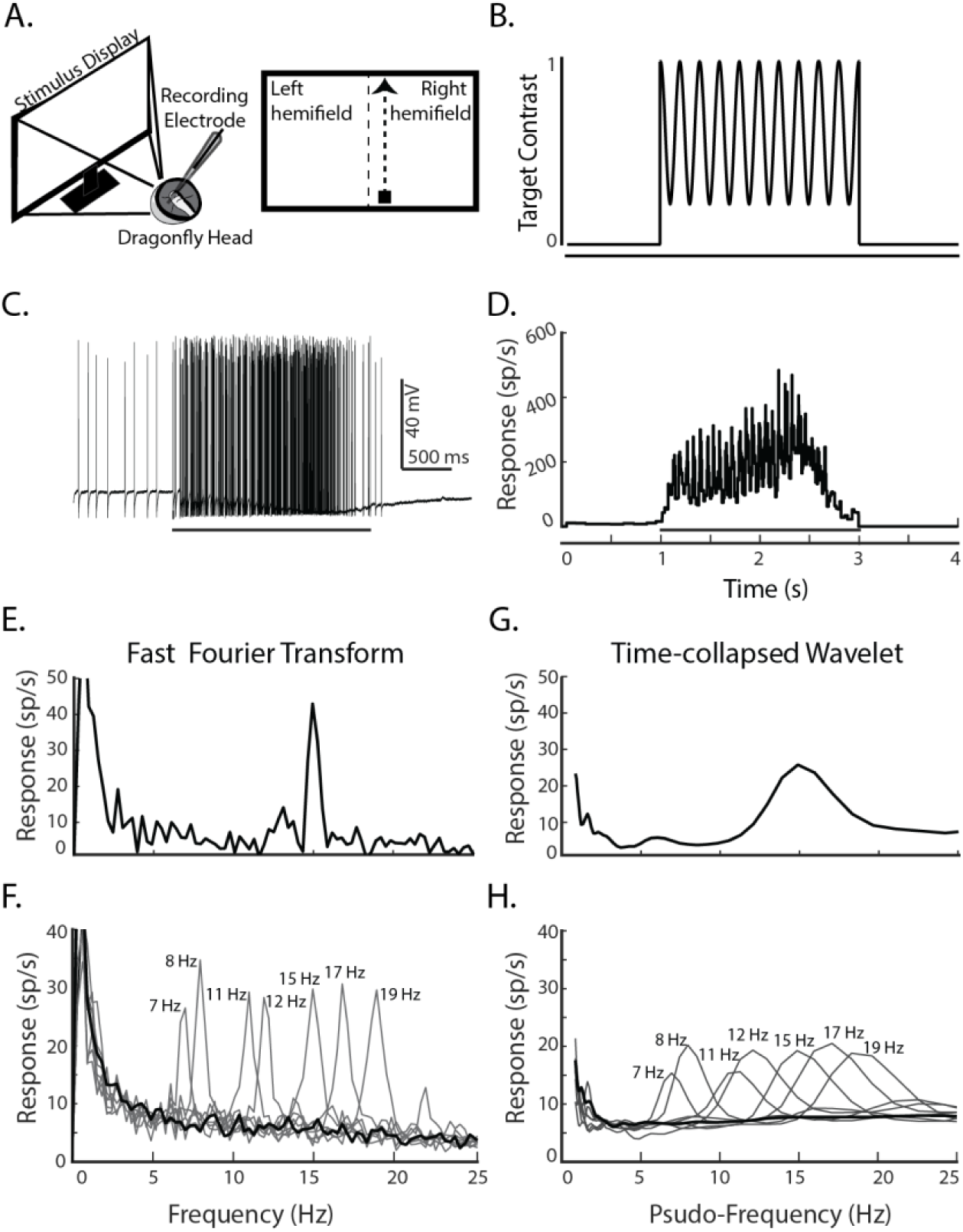
The frequency of the tagged target is preserved in the intracellular responses of CSTMD1. A) Left: intracellular in vivo electrophysiology involves inserting an electrode into the intact brain to record single-cell responses to stimuli presented on a computer screen. Right: stimulus pictogram, a single small target ascends CSTMD1’s excitatory receptive field. B) Frequency-tagging involves modulating the contrast of the stimulus over time at a specific frequency (5 Hz in this illustrative pictogram). C) An example spike train in response to a stimulus modulated at 15 Hz, presented at 1 s for a duration of 2 s (stimulus bar). D) The inverse inter-spike interval is calculated to determine the spike rate over time. This calculation provides a continuous signal that is amenable to frequency-domain analysis. E) A Fast Fourier Transform of the signal in D reveals a distinctive peak at 15 Hz, corresponding to the frequency-tagged stimulus. F) Averaged FFT of responses to trials of varying frequency (n = 119 across 4 dragonflies) G) The output of the wavelet analysis (collapsed across time to be comparable to E.) provides an alternative analysis that can preserve time-domain information (in later, non-collapsed analysis). H) Averaged time-collapsed continuous wavelet transform for the same data presented in F, which although less peaked, still reveals statistically distinctive humps at the relevant frequencies.

Can the frequency tagging technique replicate the selective attention result (Wiederman and O’Carroll, 2013) when two targets flickering at different frequencies are used? To test this, we presented either single targets (pseudo-randomly at either f_1_ or f_2_) at either spatial location T_1_ or T_2_ (both within CSTMD1’s excitatory receptive field). Randomly interleaved with the single target trials (**Figure 2A**), we also presented paired targets (simultaneously at target locations T_1_ and T_2_) which were frequency-modulated at the two *different* frequencies (pseudo-randomly between T_1_=f_1_, T_2_=f_2_ and T_1_=f_2_ and T_2_=f_1_). As our interest is in the chosen target (T_1_ or T_2_), rather than the frequency of the ‘identifier’, we pooled across the frequency-pairs.

**Figure 2:**
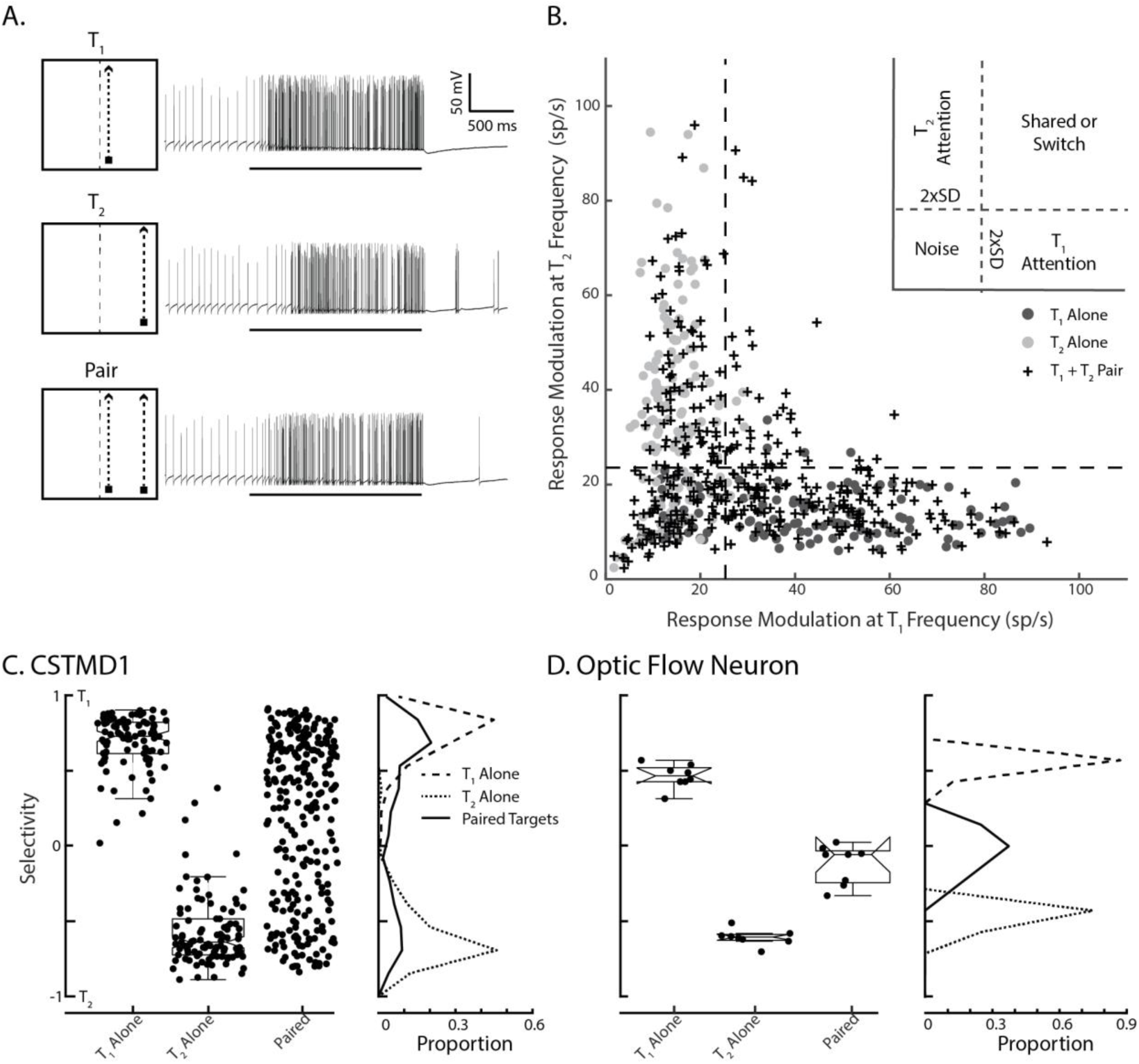
Frequency-tagging identifies the selected target in a paired-target trial. A) Illustrative pictograms and corresponding electrophysiological responses for the 3 stimulus conditions. From top-to-bottom: T1 Alone; T2 Alone; Paired Targets. B) The response modulation at the T_2_ frequency plotted against response modulation at the T_1_ frequency. Responses are plotted to a single target at the T_1_ location (dark dots) or at the T_2_ location (light dots) when presented alone. Crosses represent responses to the Paired stimulus (n = 447 trials across 13 dragonflies). Dashed lines indicate a noise threshold. Most of the responses to paired targets elicit responses at either one or other of the target flicker frequencies (not both together), indicative of a selection process C) The Selectivity index is represents the degree to which the response favours one of the frequency tagged stimuli over the other. Values around zero indicate that both frequencies are equal components of the response. Frequency polygons illustrate the relative proportion of these points, with the bimodal distribution to the paired stimulus clearly revealing the selection of one target or the other. D) In contrast, results from an optic-flow neuron in the dragonfly show no selective attention (n = 8 trials in 1 dragonfly), with a unimodal distribution around zero to the paired stimulus, indicative of shared modulation to both target frequencies.

In single target trials (**Figure 2B**, T_1_ dark dots and T_2_ light dots), we observed modulation at the frequency of the presented target and low modulation at the other frequency (i.e. a frequency that does not exist in the stimulus). However, in some individual trials there was insufficient modulation in the frequency domain to enable accurate identification of the selected targets. This is likely to result from two factors: (1) neuronal habituation in the receptive field diminishing the strength of the modulation (2) neuronal saturation from a highly responsive cell limiting the possible strength of the modulation. To analyse trials free of these effects, we used single-target responses to determine a threshold for data inclusion. For each location, T_1_ and T_2_, we calculated the average magnitude at the frequency *not presented*, which provides a value of the noise inherent in the frequency domain. This floor was defined as the mean power at the non-presented frequency plus twice the standard deviation. This provided an objective level of the modulation noise at the other frequency. That is, the expected, non-zero modulation at f_2_ when the neuron has selected a target modulated at f_1_, and vice-versa (**Figure 2B** – dashed lines). Trials in the bottom-left corner of **Figure 2B** thus fail the acceptable signal-to-noise threshold for both frequencies. 172 trials were rejected from further analysis for this reason and our frequency-target technique thus worked for 71.4% of the total trials presented. There was no significant difference in the amount of trials excluded via this metric between any of the three conditions (X^2^-test, *P > 1, Bonferroni-holm correction*). Therefore, we propose that the lack of modulation was due to ether habituation or summation, rather than related to the test under consideration – the presence of selective attention. In the successful trials, signals were above threshold at either f_1_ or f_2_, indicating significant response to either one, or both, of the targets. Qualitatively, we observe that the responses to paired targets (**Figure 2B**, crosses) were mostly either modulated at the frequency of the target at location T_1_ or T_2_ (but not both – crosses within the ‘Shared or Switch’ region).

The absolute modulation above this noise threshold (i.e. the distance of the data points along the abscissa or ordinate in **Figure 2B**) is related to the trial-by-trial sensitivity, rather than to the degree of the selective attention to one or either of the targets. To quantify our data, we therefore defined a *Selectivity Index* (**Figure 2C**), which measured the degree of target selection, independent of the strength of response modulation (though above the noise threshold previously described). For each data point, we calculated:

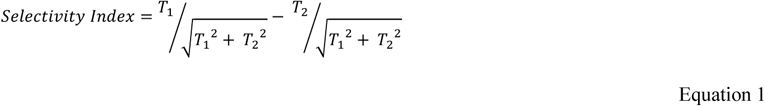

T_1_ and T_2_ values are averages of the pseudo-frequency amplitude (known as ‘scale’) over the trial duration (i.e. collapsed across time from the CWTs), for each of the corresponding target frequency-tagging modulations. The selectivity index ranges between +1 and −1 and represents the selection of T_1_ (+1) and T_2_ (−1), respectively. Here ‘selectivity’ is referred to in the original definition of ‘selective attention’ as selection of one from multiple competing stimuli, as would be expected in a winner-takes-all network. A value of 0 would occur if the response magnitude at f_1_ and f_2_ were equal (irrespective of the absolute distance from the origin), indicating either shared (co-varying) selection across the trial, or a switch in selection during the trial.

In **Figure 2C**, we observe significant differences in the Selectivity Index distribution between paired and both T_1_-alone and T_2_-alone conditions (P < 0.001). In single-target conditions, the Selectivity Index is narrowly distributed (T_1_ µ = 0.68, σ = 0.17; T_2_ µ = −0.58, σ = 0.23), whereas in paired-target trials the Selectivity Index is non-normal (p > 0.001, one-tailed Kolmogrov-Smirnov test) with peaks at approximately 0.65 and −0.55. The bimodal distribution of responses to paired targets reveals the selection of either T_1_ or T_2_. For comparison to a potential ‘null’ hypothesis (i.e. no selective attention), **Figure 2D** shows results from a single ‘optic flow’ neuron in the dragonfly. This neuron generates robust responses using spatial summation in order to encode wide-field optic flow, analogous to Lobula Plate Tangential Cells in Diptera (Hausen, 1982). We presented the same experimental paradigm, though with larger targets (1.5° x 10°) to elicit a response. In contrast to the results observed in CSTMD1, the optic-flow neuron had a Selectivity Index around 0 (modulation at both frequencies of the paired targets) indicative of neuronal spatial summation.

Not all of the paired-target trials were solely modulated by one of the target frequencies (**Figure 2B**, shared zone). If CSTMD1 is only selectively attending one of the presented targets, what could account for this apparent shared modulation? There are two possible explanations: firstly, the neuron is excited by both stimuli at their respective frequencies and not selecting a single target. That is, spatial summation similar to what is observed in the Optic Flow neuron (**Figure 2D**) and in primate V4 (Ghose & Maunsell, 2008). Alternately, a switch mid-way through the trial could result in significant modulation at both frequencies, as both targets are selected during the trial, though at discrete times.

To test this possibility, we first simulated a switch in response from f_1_ to f_2_ by presenting a single-target that changed frequency in the middle of the trial (**Figure 3A**). An example of the intracellular response to such a pseudo-switch stimulus is presented in **Figure 3B**. We subtracted the two wavelet magnitude ‘slices’ from one another (**Figure 3C**, dashed lines) derived from the CWT analysis, thus producing a difference in magnitude between the two pseudo-frequencies over time (**Figure 3D**). This difference in magnitude highlights the degree to which one frequency is selected over the other throughout the time course. For example, capturing the frequency change in in the simulated switch with a peak and trough dissociated in time. This technique therefore provides a ‘read-out’ through time of when the response was governed by which frequency. In a case where power was shared, we would expect a flat line representing little variation in the frequency magnitudes across time.

**Figure 3:**
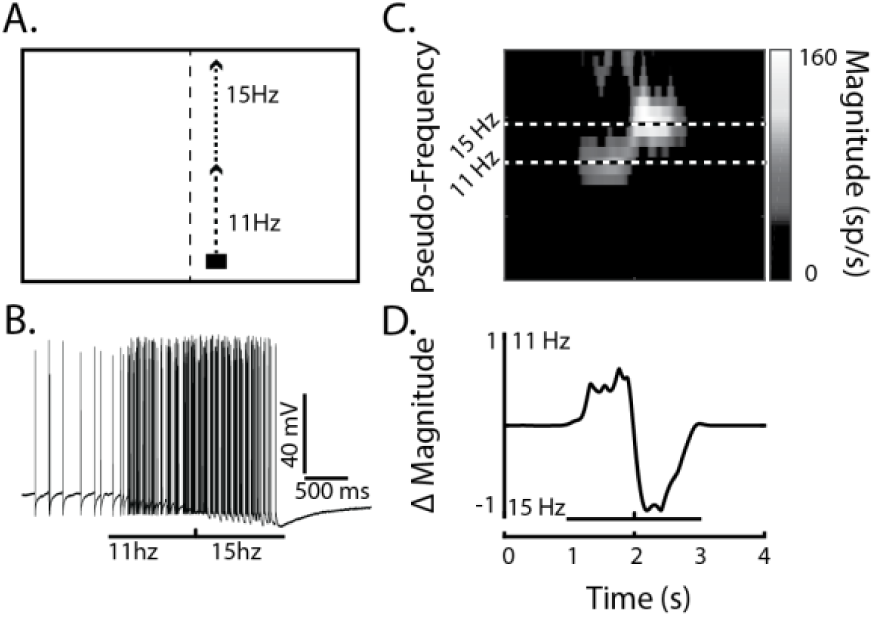
Simulation of an ‘attentional switch’ produces a characteristic result. A) Illustrated pictogram of a single target that changed frequency halfway through the trial, simulating an attentional switch from a target of one frequency to the other. B) An example of a CSTMD1 response to this switching stimulus. C) The CWT analysis of the inverse ISI of the trial in B, reveals the switch that occurs halfway through the trial. The black-and-white dashed lines indicate the 11 Hz and 15 Hz frequency slices. D) A ‘difference slice’ (delta magnitude) is calculated by taking the difference between the wavelet slices at 11 and 15Hz across time.

We applied this ‘read-out’ analysis to determine whether the paired target responses with modulation at both frequencies (**Figure 2B**, shared or switch region) were due to spatial summation or switching. **Figure 4A** shows examples from six such trials, all of which exhibit discrete peaks and troughs in time. The traces indicate that these CSTMD1 responses are switching between targets over time, rather than being modulated by both target frequencies simultaneously.

**Figure 4:**
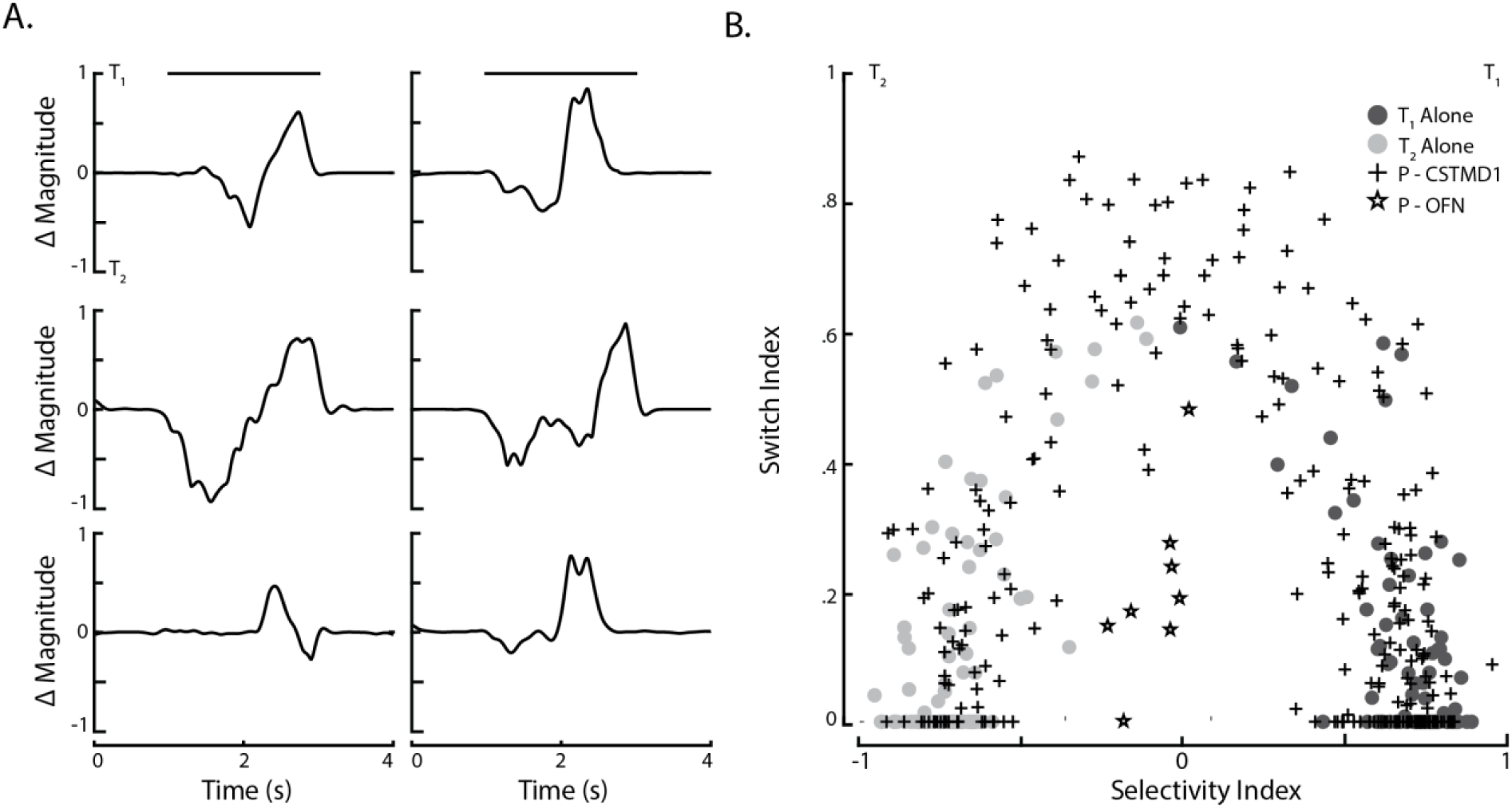
Shared modulation results from switches in selection. A) Individual trial examples of the ‘difference slice’ from the wavelet analysis of paired-target trials showing high modulation for both targets at different epochs of time. B) The ‘Switch index’ and ‘Selectivity index’ for all single target (dark and light points) and paired target (crosses) trials. When selectivity for paired targets is low (middle abscissa, close to zero) then the Switch Index is high, indicating that responses switched between targets. Instead, the optic flow neuron (stars) has low Selectivity and a low Switch, indicative of response summation (modulation at both frequencies across points in time).

To compare aggregate data, we calculated a ‘Switch Index’ for each trial (**Figure 4B**). This index was calculated by first determining the proportion of time the system selected either T_1_ or T_2_. To ensure that these selections were robust we only considered a selection valid when either target was significantly stronger (> 5 spikes/s) than its counterpart (i.e. when T_1_ > T_2_ + 5 or vice-versa). Having established how long each target was selected, we then multiplied these two values together. This has the effect that when one of the two targets was not selected at all, the Switch Index is zero, while it is maximized when both targets are selected for 50% of the trial. This value was normalized between 0 and 1. In single-target trials (dark or light dots), the Switch Index is overall low, however in paired-target trials the Switch Index is distributed between high and low values. In trials with a Selectivity Index around 0, the Switch Index is uniformly high, indicating that the low selectivity is almost entirely due to switches. In contrast, paired-target trials in the dragonfly optic flow neuron show both a low Selectivity Index and low Switch Index, indicating genuine modulation at both frequencies over time due to the spatial summation used in optic flow computations (**Figure 4B**, stars).

### Biasing selection with priming

Next, we tested the ability of a priming stimulus to *bias* the selection of a spatially-associated target in a paired-target condition. In this experiment, a lone untagged primer was first presented for one second moving towards the trajectory of either spatial location T_1_ or T_2_ (**Figure 5A**). Note that here the frequency-tagged T_1_ and T_2_ pathways commence midway up the stimulus display, immediately after the single ‘primer’ target has moved along its trajectory. From our previous work, we expect CSTMD1 to ‘lock-on’ and predictively facilitate responses in front of the target’s prior path (Nordstrom 2011; Dunbier et al, 2012; Wiederman & Fabian et al., 2017). In this experiment, we introduced a frequency-tagged distracter midway through the receptive field (horizontally offset by 20°) paired with a frequency-tagged target that continued along the previous ‘primer’ targets’ trajectory (**Figure 5A**). We calculated the Selectivity Index across the entire second where both targets are presented together and reveal a significant (*P < 0.001*) biasing of selection towards the target that continues along the primed trajectory (**Figure 5B**). This selection may be attributed to the previously described predictive gain modulation, whereby a local spotlight of enhanced gain is generated ahead of a moving target, with suppression in the surround (Wiederman & Fabian et al., 2017). In our experiment, the continuing target is within the spotlight created by the preceding target, but the distracter appears within the supressed surround.

**Figure 5:**
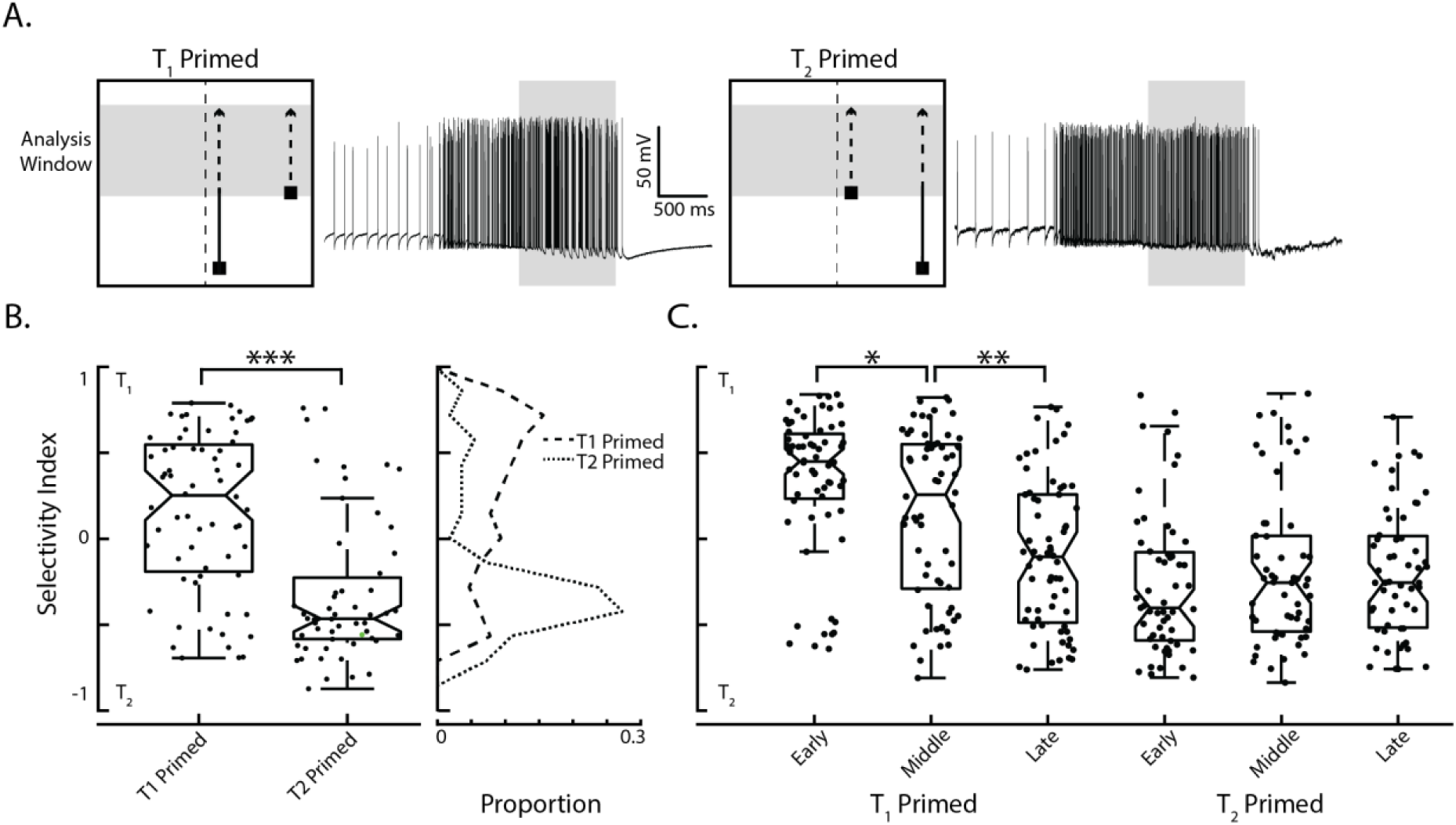
Priming with a preceding target biases selection towards the continuing trajectory. A) Pictograms illustrate the biasing stimulus towards either spatial location T_1_ or T_2_, next to individual example of CSTMD1 responses to the stimuli. The short-path target (distracter) appears at 1 second, when the preceding target reaches midway up the screen (the analysis window indicated with the grey shade). B) There is a significant difference between the Selectivity Index when T_1_ was primed with the preceding target compared to when T_2_ was primed (n = 295 across 7 dragonflies). Frequency polygons reveal the distributions of the Selectivity Index for T_1_ primed (dashed line) and T_2_ primed (dotted line). For comparison, we also plot the previously described paired target data (solid line) of the two frequency-tagged targets moving along the entire T_1_ and T_2_ pathways C) In order to assess the impact of attentional capture we split the paired target period into 3 windows which were analysed separately.

In the human psychophysics literature, attentional capture is an effect whereby the presentation of an abrupt-onset stimulus (Yantis & Jonides, 1984) or a novel object (Franconeri, Hollingworth & Simons, 2005) involuntarily captures attention (Remington, Johnston & Yantis, 1992), even when task-irrelevant. In order to test for a capture of CSTMD1’s selection, we analysed the previous biased paired-target responses (**Figure 5B**) separated into three 400 ms periods (Early, Middle and Late). We included 100 ms overlap between these periods because this duration was required for meaningful CWT analysis. If CSTMD1 responses displayed attentional capture, we hypothesise that the early period would be dominated by responses to the distracter stimulus, returning to the original path at later periods of time (as the distracter is assessed and ignored). Our results reveal the opposite effect (**Figure 5C**), with the early window exhibiting the strongest effect of the biasing which dissipates over time. This reveals that the selection is not automatically captured by the abrupt-onset novel stimuli presented within CSTMD1’s receptive field, rather responses are locked on to the preceding target. Here we observed asymmetry in results from the T_1_ compared to T_2_ priming, which reflects the broader (noisier) distribution of values in the T_1_ Primed condition when analysed over the entire analysis duration (**Figure 5B**). When primed to T_1_ (the target closer to the dragonflies’ midline), the Early window (**Figure 5C**) reflects this biasing to the continued path trajectory (though note the several clear exceptions). However, in some cases over time (Middle and Late windows) selection changes towards the distracter location at T_2_, resulting in significant changes in the Selectivity Index between these periods (*P < 0.001*). Note that visual inspection of the CWT analysis reveals that these are switches that occur at specific points in time in the individual trials. In the T_2_ priming condition (the target located in the more peripheral location), the selection has locked on to the preceding target and maintains this selection throughout the rest of the trial, with no significant difference between the Early, Middle and Late periods (again with several clear exceptions).

In a traditional winner-takes-all network, the introduction of a high contrast distracter during the presentation of a lower contract target would result in a switch to the one with higher salience. However, how would the dragonfly feed in a swarm if often distracted by a novel, transiently more salient target? To determine whether CSTMD1 locks-on to the lower-salience stimuli, we presented primers of varying contrasts followed by a paired frequency tagged distracter. In this experiment, we wanted the lower contrast target to retain its lower saliency throughout the course of the trial, even during the period when the frequency-tagged distracter was present. Because the original target (of varying contrast) is never frequency-tagged, it is the measure of modulation that is the indicator of distracter selection. Primers were presented at constant Low (0.06), Medium (0.15) or High (0.51) contrast, pseudo-randomly located at spatial locations T_1_ or T_2_ (**Figure 6A**, primer at T_2_ location shown). The High contrast primer was set at 0.51 to be equiluminant with the *average* contrast (over time) of the frequency-tagged, high contrast distracter. **Figure 6A** shows example responses of an individual CSTMD1 to these stimulus conditions, both to when the primer retains selection and when the distracter takes over. This shows that there can be trial-by-trial variability in which of the targets was selected (primer or distracter).

**Figure 6:**
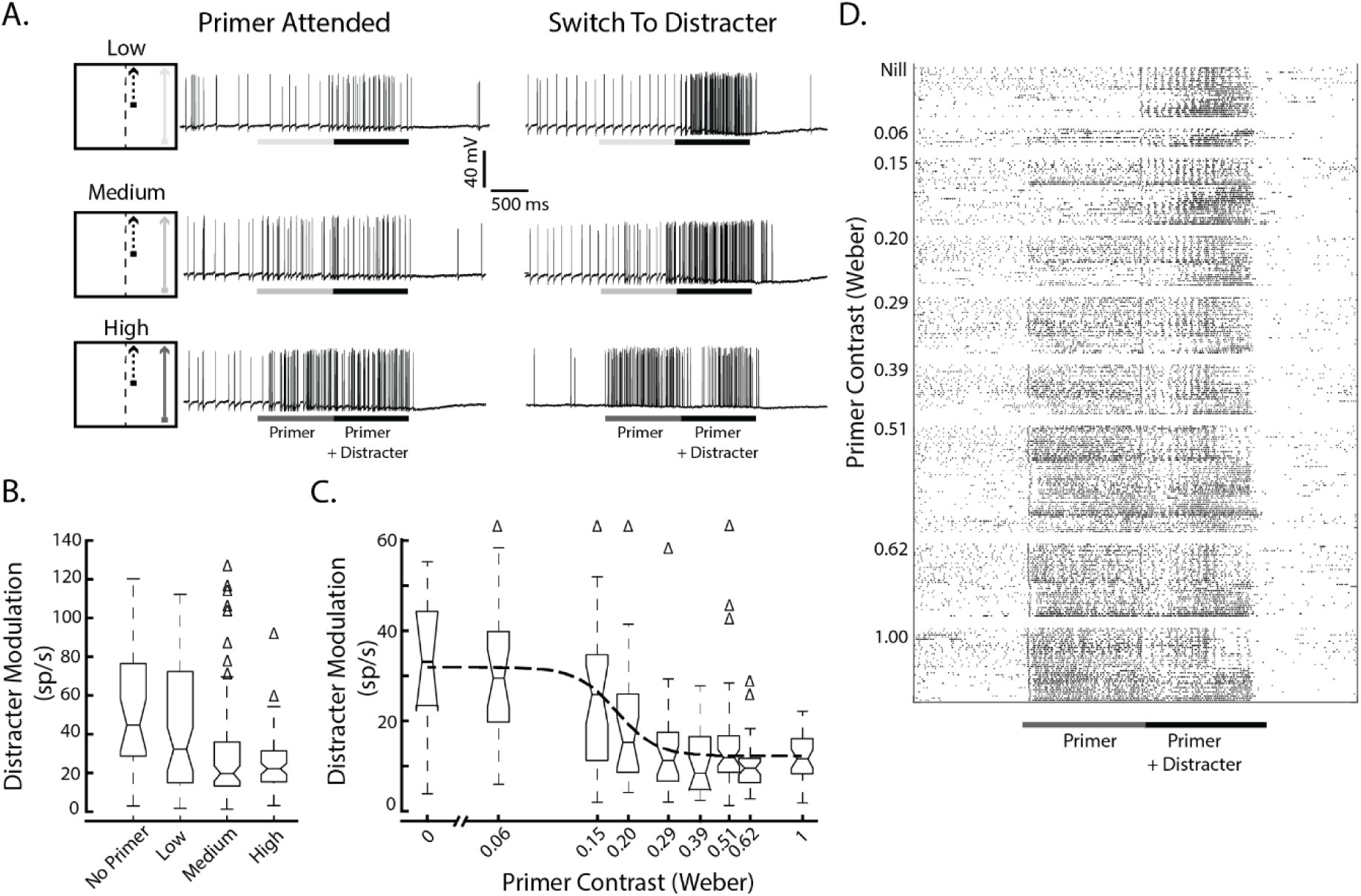
Selective attention in CSTMD1 can ‘lock on’ to a lower contrast target, ignoring the introduction of a high contrast distracter (frequency-tagged. (A) Stimulus pictograms and example traces from the same CSTMD1 for Low, Medium, and High contrast primer conditions. B) Boxplots showing the modulation at the distracter frequency across four primer conditions (n = 204 across 5 dragonflies). Δ indicate outliers. The distracter only condition (No Primer) shows the expected distribution of modulation values if the priming target is never selected. Even in the Low contrast condition, there is a significant difference, indicative that in some trials the Low contrast target is selected during the period when a high contrast distracter is present. As the salience of the primer is increased, the number of trails where the distracter is selected decreases C) In an individual CSTMD1 recording, we assayed across a larger range of primer contrasts, revealing a sigmoidal contrast sensitivity function (n = 212 across 1 dragonfly). Δ indicate outliers. Off-axis outliers are indicated. D) Spike rasters organized by primer contrast from the individual CSTMD1 data presented in C. with trials within each contrast are ordered by presentation time.

**Figure 6B** shows a significant reduction in distractor modulation as primer contrast rises (Kruskal-Wallis One-way ANOVA, df = 3, *χ*^*^2^*^ = 21.32, p < 0.001), indicating that CSTMD1 can lock-on to Low contrast targets even in the presence of a high contrast distracter. There is still trial-by-trial variability, however as primer contrast increases a higher proportion of trials do not exhibit distracter modulation, thus have selected the primer. Due to the previously described biasing effect of a preceding primer (**Figure 5**), we would expect less distracter modulation at the equiluminant contrast (higher than 0.51). We observed a significant reduction in distracter modulation in the medium (P =.006) and high (P >.001) contrast group, but not the low-contrast group (P =.755), compared to the no primer group (**Figure 6B**), suggesting that CSTMD1 was indeed able to lock on to primed targets of.15 and higher contrast. However, outliers observed in both the Medium and High contrast conditions indicate that CSTMD1 is still able to switch selection to the distracter on some trials, consistent with the previously observed “rare” switching which occurred with two equally salient targets (Wiederman & O’Carroll, 2013).

From a single CSTMD1 recording, we were able to assay across more primer contrasts (**Figure 6C**). In this neuron, the primer often locked-on to the much lower primer contrasts (as low as 0.2), ignoring the simultaneously presented distracter target. Therefore the mechanism underling this neuronal selective attention cannot be a ‘simple’ winner-takes-all network. Raster plots of spikes throughout the trial reveal some interesting attributes of the response (**Figure 6D**). Even in trials where the distracter was not selected, the onset was marked with a reliable spike, perhaps indicative of a transient breakthrough in the underlying network. Additionally, in conditions when the primer contrast is higher, there is often a transient suppression of response (timed with the distracter onset) before responses continue with their selection. Further experiments are required to elucidate how this phenomenon relates to the suppressive surround observed in the predictive gain modulation observed to single target trajectories (Wiederman, Fabian et al. 2017)

Overall, these data clearly reveal that CSTMD1 is able to lock on to low-contrast targets and select them even in the presence of a high contrast distracter. Intriguingly, response to these continued primer trajectories are not associated with an increase in spike rate as would be expected by models of attention where low-contrast stimuli are attended by neuronally boosting contrast (Reynolds & Desimone, 2003). Instead, even when responding to low-contrast stimuli, CSTMD1 encodes the absolute strength of the attended target as if the distracter was not present (**Figure 6A**). This could be critically important in behaviour where a target is selected for pursuit amidst a swarm, where absolute rather than relative activity might underlie the closed-loop control system.

### Modelling

What mechanism best explains the measured data? To test this, we developed six algorithmic models. The six models included two models that assumed shared attention (including one with saturation), two models that applied selection and two models which applied selection with switching. For input to these models we collected the single target trial (i.e. T_1_-only or T_2_-only) response modulation amplitude from the wavelet analysis (**Figure 2**).

From this we produced four lists (T_1_f_1_, T_1_f_2_, T_2_f_1_, T_2_f_2_) representing the response modulation amplitude at the target’s flicker frequency and at the comparison frequency (i.e. no modulation). We binned these responses and fit a log-normal distribution to each target and frequency pair (T2 examples are shown in **Figure 7A**). We were then able to infinitely sample from these model distributions to generate an arbitrary number of synthetic target responses.

**Figure 7:**
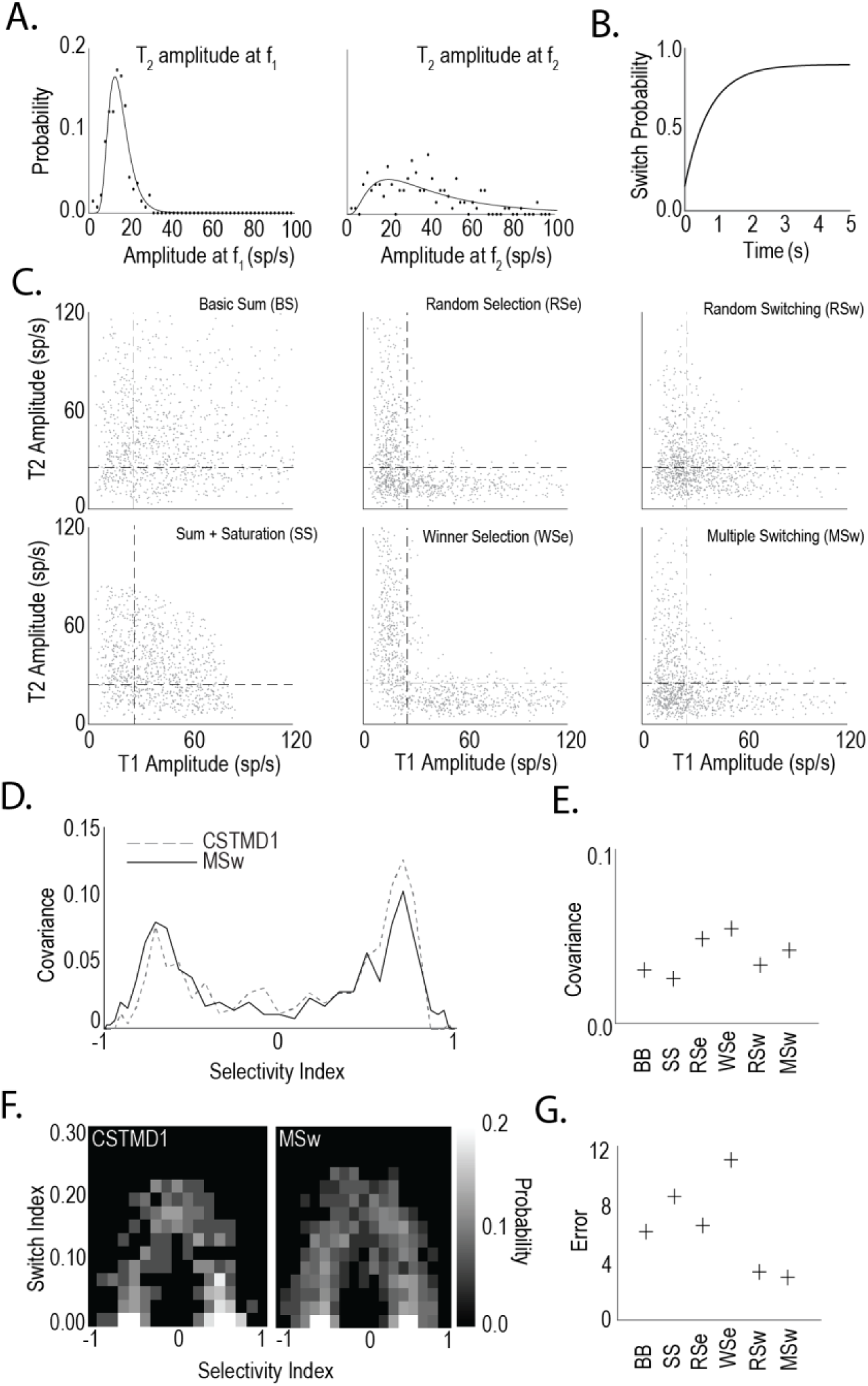
Simple Switching Models Matches Selective Attention Data. A) Power distributions for frequency responses from T_2_ at f_1_ (left) and f_2_ (right) calculated from recorded trials. Modelled trial data were randomly selected from these power distributions representing the power contribution of each target. B) Switch probability as time progresses for model 6 (Multiple Switching). Initially the likelihood of switching is low before rising to 90%. After a switch, the switch probability resets allowing multiple switches to occur. C) Example scatter plots (as per **Figure 2**B) for each of the six models tested. Summation (top left), summation with saturation (bottom left), random selection (top middle), higher power always wins selection (bottom middle), random switching (top right), and multiple switch model (bottom right). D) Histogram of model selectivity for recorded data and model output. Error calculation as covariance curve. E) Results of six models against recorded data from histogram analysis shown in (d). Higher covariance is indicative of a more representative model. F) Two-dimensional histogram (selectivity/switch index) for recorded data (left) and model data (right). Error calculated as RMS deviations from recorded data. G) Results of six models against recorded data using 2D histogram data (f). Low values indicate representative model.

Simulating switches requires a time-course of the response modulation amplitudes over time. To simulate this, we generated a 1s time course for testing all models (even non-switching models). These time-courses represent the instantaneous response modulation amplitude over time equivalent to taking a 1-dimensional slice from the CWT analysis (as in **Figure 3**). It is unrealistic that the response modulation amplitude at a given frequency would be constant over a 1s period. To ensure a more realistic response, we added noise (white noise with a 5mV max width). We then smoothed the data using a 0.2s average filter which produced waveforms qualitatively similar to those observed from taking a single-frequency slice of a CWT. For switching models the smoothing was done after calculating the switch. For example, if a switch from T_1_ to T_2_ occurred at 0.4s, the first 0.4s would use both the T_1_f_1_ and T_1_f_2_ (each with noise added) while the subsequent 0.6s would use T_2_f_1_ and T_2_f_2_ (again with noise added). This process regularly produced ‘step-like’ responses to which we applied smoothing (as mentioned above) to generate the smooth transitions we saw in the CWT data.

We sampled from the distribution 1000 times for each pairing (T_1_f_1_, T_1_f_2_, T_2_f_1_, T_2_f_2_). Each model used some combination of these to generate output responses for both f_1_ and f_2_. Basic Summation (BS) assumed that the output power at both f_1_ and f_2_ were the corresponding powers of the input target (i.e. T_1_f_1_ & T_2_f_2_). Saturating Summation (SS) summed like BS, but applied a soft saturation to reduce the overall modulation power evenly between f_1_ and f_2_ to a maximum potential power of 100 spikes/s. Random Selection (RSe) randomly selected either T_1_ or T_2_ and used that target’s corresponding power for f_1_ and f_2_ (i.e. if T_1_ was selected the frequency responses would be T_1_f_1_ and T_1_f_2_). Winner Selection (WSe) selected the target with the greatest modulated power (using the tacit assumption that the modulation was proportional to the target response) and used the winner’s frequency response solely. Thus if T_1_f_1_ > T_2_f_2_, T_1_ would be selected and vice versa. Random Switching (RSw), randomly selected an initial target (as per RSe) but assumed that a switch occurred in a percentage of trials at some point during the trial’s duration (specific values determined after optimization, see below). Multiple Switching (MSw) assumed a more sophisticated switching rate, allowing the system to switch multiple times. The switch probability was defined by the following formula:

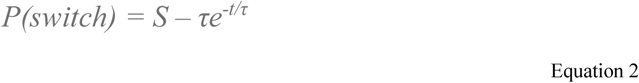

S represents the probability that a switch never occurs and τ represents the rate of increase of switching over time (**Figure 7B**). The values of S and τ chosen were determined after an optimization step (see below).

The generated outputs of all six models are shown in **Figure 7C**. The summation model (BS) populates all four quadrants (including in the ‘Shared or Switch’ zone of Figure 2B). This combination of taking power from both targets together does not match the electrophysiological results (**Figure 2B**). Both selection models (RSe & WSe) adhere far closer to the distribution seen in **Figure 2** except that the shared zone is very sparsely populated, especially in the WSe model. The switching models are good qualitative matches for the real data with a bias to T_1_/T_2_ only responses (the L shape) but with a reasonable number of shared zone responses indicative of switching.

To assess each model quantitatively, we generated the frequency polygon (**Figure 2**, **Figure 4**) of the selectivity index values calculated from the model outputs. An example of the response of the MSw model (grey line) compared to the electrophysiological data (dotted line) is shown in Figure 7D. We compared each model’s frequency polygon with frequency polygon from **Figure 2C** via cross-correlation.

Via this metric, both selection models (RSe, MSe) provided the best match to the recorded data (**Figure 7E**). However, the selection metric ignores the switching behaviour inherent in the model. To test whether pure selection was sufficient to explain the data, we used the model outputs to calculate the ‘Switch Index’ (**Figure 4**) for each model’s responses. We then binned this data to generate a 2-dimensional histogram (**Figure 7F**). We repeated this process for the electrophysiological data and calculated the RMS error between the two. As both switching models had free parameters (i.e. probability of switching) we optimized both these models against this RMS error. The RSw model was most successful with a 100% probability of a switch at a random time during the trial. The MSw model was optimal with a 90% switch probability and 0.75s time constant. The remaining models (Summation and Selection) did not have any parameters to effectively optimize.

**Figure 7G** shows the results of the Switch Index comparison for the six models. Both switching models have lower RMS than the other models with the multi-switch model performing the best overall. It is clear from both the qualitative and quantitative aspects that the switching models produce the best outcomes. This is in line with our expectations. As mentioned previously, the Summation models (BS, SS) generate too many responses in the shared zone by effectively increasing the overall power, while the Selection models (RSe, WSe) go the opposite direction effectively eliminating most of the responses from the shared zone. The Switching models (RSw, MSw) provide a suitable compromise, with a general shift towards the upper-left and lower-right quadrants while maintaining some responses in the shared zone, However, while there were numerous responses in this region they have lower overall power indicative of temporal sharing (rather than summation). It is clear that the best explanation for the results seen is a model that selects a single target but is capable of switching one or more times during a trial.

## Discussion

Frequency-tagging techniques have previously been used during higher-order brain measurements (e.g. EEG) or in extracellular recordings measuring local field potentials (LFP) in insects (van Swinderen, 2012). However, it is not yet known whether frequency components within the frequency-tagged LFPs originate at the level of single neurons, or are an emergent property of a neuronal population code. To our knowledge, here we present the first application of this identification technique at the intracellular level. We thus demonstrate that the frequency component of the stimulus is preserved in the response of an individual neuron.

Frequency tagging allows us to verify previous findings of selective attention in CSTMD1 (Wiederman and O’Carroll, 2013) and for the first time identify which of a pair of targets was selected at any moment in time. However, it is clear that frequency tagging is not always robust. In approximately 25% of trials, regardless of stimulus conditions, levels of frequency modulation were below-noise despite a reasonable spiking response (**Figure 2B**; bottom-left corner). These trials were excluded as identification of the selected target could not be achieved. Difficulty in choosing the correct stimulus waveform may underlie this problem: Firstly, flickering targets located within the strongest parts of the receptive field may saturate, resulting in a lack of headroom for significant modulation. Conversely, frequency-tagged targets presented in less sensitive regions of the receptive field may not elicit responses strong enough to carry modulation over the underlying signal. Both factors, saturation and sensitivity, can vary dynamically as overall CSTMD1 responsiveness may change over time, location or between animals. These effects could be minimized by changes to the stimulus waveform, with a lower mean level of intensity accounting for saturation and a higher amplitude of contrast modulation for sensitivity. However, as these exclusions did not affect our hypothesis testing (were distributed equally across all experimental conditions), we kept the amplitude and mean level consistent across all experiments.

Although frequency tagging was used as an identifier, could the frequency itself interact with facilitatory or selective processing? Such a factor can play a role in other animal models, with honeybees preferencing 20-25 Hz and avoiding 2-4 Hz visual flicker (Van De Poll et al., 2015). Even a single luminance change is enough to break inattentional blindness in humans (Palmer et al, 2018). To minimise this possibility, we distributed the two tagging frequencies across the two spatial locations (T_1_ and T_2_) as well as testing our entire experimental paradigm at two different frequency-tagged pairs. Throughout these experiments, we did not observe any effect of the frequency-tagging beyond our intended purpose as an identification technique.

Attention is a limited resource (Alvarez & Franconeri, 2007), therefore animals across species are motivated to guide the deployment of attention in an ethologically meaningful and efficient way. One guide is spatial or temporal cueing, often through inhibitory neural mechanisms (Romer et al., 2002; Ruthruff & Gaspelin, 2018). For example, Drosophila are more likely to orient towards cued locations of the receptive field when subsequently presented with multiple targets (Sareen, Wolf, & Hisenburg, 2011). Female crickets prefer leading male auditory signals to signals arriving later (Snedden & Greenfield, 1998; Romer, Hedwig & Ott, 2002), suggesting an inherent bias towards ‘locking on’ to the first stimulus and ignoring those subsequent. This is similar to what we have observed in CSTMD1, with the priming by a preceding target biasing selection to those that continue along the projected trajectory.

In CSTMD1, the effect of spatiotemporal cueing was so strong that even targets of lower visual salience can win over the simultaneously presented distracter. In attentional networks, saliency is a prominent attribute for guiding selection and seems to innately capture attention. This leads to a conundrum; if the most salient objects were to capture attention moment-to-moment, then the system might too often be distracted from any given task. For example, will the dragonfly ever feed if the prey of varying contrast (i.e. moving against a cluttered background) becomes less salient than others in the swarm? Conversely, the onset of a novel salient stimulus may signal the necessity to attend to a new event or abandon the current task completely in favour of survival behaviour (e.g. an approaching bird).

In human psychophysics, both abrupt-onset (Yantis & Jonides, 1984) and perceptually new objects (Franconeri, Hollingworth & Simons, 2005) provoke attentional capture, a phenomena where attention is automatically and involuntarily directed at a particular, often task irrelevant, feature (Remington, Johnston & Yantis, 1992). The signal suppression hypothesis by Sawaki and Luck (2010) proposes that all stimuli automatically generate a saliency signal, but this signal can be supressed by top-down attentional mechanisms. In our CSTMD1 recordings, we found no evidence for attentional capture. Instead, the earliest period of the paired targets revealed the strongest bias to the previous primer trajectory, with the possibility of switching to the more novel distracter at a later time. Thus rather than attending to a novel distracter, this system is locking on to the expected target trajectory. These results may be attributed to the previously observed effect that CSTMD1 predicts future target location following an occlusion (Wiederman, Fabian et al, 2017) with an enhancement for the prior path and suppression in the surround. During the initial window, the continuing target is fully facilitated by the preceding target and continuously moving into its self-generated spotlight of gain enhancement. However, the distracter appears within the supressed surround and therefore will not elicit attentional capture (Ruthruff & Gaspelin, 2018) in agreement with the signal suppression hypothesis (Sawaki & Luck, 2010). By the middle period, the distracter may have self-facilitated, enabling a more even competition for target selection and thus increasing the probability of a switch. Whether this self-facilitation occurs at both target locations before selection, or only at the single selected location is currently under investigation.

These results bear resemblance to behavioural results in *Drosophila* (Koenig, Wolf & Heisenberg, 2016). Tethered flies in an arena were presented with a pair of vertical lines equally offset from the flies’ midline. Flies made a decision to respond to either one line or the other by turning to bring it into the midline. In subsequent trials, these flies displayed a bias for turning towards the originally selected stimulus and ignoring the alternative. However, over time this bias was lost. The mean ‘attention span’ (time before the bias was lost) was 4 seconds in wild-type flies, but reduced to 1 second in mutants defective in selective attention. Active switching between competing stimuli may be indicative of endogenous drive by top-down control mechanisms (Miller, Ngo & van Swinderen, 2012). Van Swinderen (2007) found that, in Drosophila, a minimum amount of time must pass between the original selection of a target and switching to a new stimulus, and switching at all was reliant on short-term memory genes.

The possibility that non-selected stimuli also generate a spotlight of neuronal gain modulation is in agreement with proposed mechanisms underlying attention in primates (Reynolds & Desimone, 2003). Primate cortical cells are thought to be ‘hard-wired’ to respond to the highest contrast stimulus, a property that can be exploited by attentional systems in V4 (Schiller & Lee, 1991; DeWeerd et al., 1999). Here the representation of stimuli is modulated by enhancing the effective contrast of the focus of attention (Martinez-Trujillo & Treue, 2002; Reynolds & Desimone, 2003). Through this enhancement, less salient and even non-preferred stimuli can come to dominate the response of neurons in V4 (Reynolds & Desimone, 2003), MT, and MST (Recanzone, Wurtz & Schwarz, 1997; Treue & Maunsell, 1999).

This neuronal enhancement observed in primates may be mechanistically similar to the facilitation observed in CSTMD1, where in response to a single target gain is increased ahead of the prior path and suppressed in the surround. In primates it is the presence of distracters that triggers this attentional enhancement (Treue & Maunsell, 1999; Reynolds, Pasternak & Desimone, 2000; Treue, 2001; Reynolds & Dismone, 2003). However, In CSTMD1, facilitation enhances the neuronal response to even an individually presented single target. In the presence of distracters the facilitated strength of the selected target is retained as if the distractor did not exist.

The ability of a neuron to respond with the same strength to a target presented alone, or when selected in a pair, may underlie the dragonflies’ exceptional ability to hunt in swarms (Combes et al, 2012). Such neuronal processing may have evolved overcome the confusion effect by singling-out targeted prey amidst a swarm (Landeau & Terborgh, 1986). Behavioural studies in some dragonfly species, *Libellula* adults (Combes et al, 2012) and nymphs (Jeschke & Tollrian, 2007) show that they are adept at hunting in swarms throughout life. Although not tested in *Hemicordulia*, this hawking dragonfly is also likely to benefit from neuronal processing that reduces the confusion effect via selective attention, as they spend most of their adult life hunting and patrolling territory on the wing including in swarms or prey and conspecifics.

